# Decoding protein-peptide interactions using a large, target-agnostic yeast surface display library

**DOI:** 10.1101/2025.05.19.654863

**Authors:** Joseph D. Hurley, Irina Shlosman, Megha Lakshminarayan, Ziyuan Zhao, Hong Yue, Radosław P. Nowak, Eric S. Fischer, Andrew C. Kruse

## Abstract

Protein-peptide interactions underlie key biological processes and are commonly utilized in biomedical research and therapeutic discovery. It is often desirable to identify peptide sequence properties that confer high-affinity binding to a target protein. However, common approaches to such characterization are typically low throughput and only sample regions of sequence space near an initial hit. To overcome these challenges, we built a yeast surface displayed library representing ∼6.1 × 109 unique peptides. We then performed screens against diverse protein targets, including two antibodies, an E3 ubiquitin ligase, and an essential membrane-bound bacterial enzyme. In each case, we observed motifs that appear to drive peptide binding and we identified multiple novel, high-affinity clones. These results highlight the library’s utility as a robust and versatile tool for discovering peptide ligands and for characterizing protein-peptide binding interactions more generally. To enable further studies, we will make the library freely available upon request.

## Introduction

Interactions between proteins and peptides underlie fundamental aspects of biology. Specifically, the ability of proteins to selectively recognize and bind peptides or short, unstructured regions of proteins is critical for numerous biological functions^1^. This includes proteins involved in adaptive immunity (e.g., antibodies^2^ and major histocompatibility complexes^3^), proteostasis (e.g., E3 ubiquitin ligases^4,5^ and heat shock proteins^6,7^), and transmembrane signal transduction (e.g., G protein coupled receptors^8^, receptor tyrosine kinases^9,10^, and integrins^11–13^). As a result, peptides are widely used as research tools and comprise an important class of therapeutics, with more than 80 FDA-approved peptide drugs to date^14,15^.

Due to their biological significance, straightforward synthesis (including biosynthesis), and success as therapeutics, peptides are commonly screened in basic research and drug discovery efforts. In some types of screens, such as mutational scans^16–19^, structure guided mutagenesis^20,21^, or antibody epitope mapping^22^, an initial hit is already known and it is sufficient to test a relatively small number of variant peptides. In these cases, individual peptides can be readily synthesized via solid-phase chemistry^23,24^ or encoded in a defined oligonucleotide pool^25^. However, these approaches are prohibitively expensive and time-consuming at scales greater than hundreds or thousands of peptides.

In cases where an initial hit is not available, broader screening must be undertaken using a larger and less biased peptide library. Such libraries have been developed in various display systems, such as phage^26,27^, yeast^28,29^, mRNA^30–32^, ribosomal^33,34^ and bacterial^35–37^ display. Although some of these library formats are theoretically capable of harboring immense diversity (≥10^14^ unique clones), such diversities are not commonly achieved in practice and libraries are typically still modest in size when compared to peptide sequence space^32,38^. Additionally, some of these libraries are designed to contain specific sequences (e.g., those found in the human proteome^27^) and are thus intentionally limited in the types of hits they can provide. Lastly, most of these library formats are incompatible with Fluorescence-Activated Cell Sorting (FACS), limiting their practical use in screening for properties beyond simply affinity, such as pH dependence^39^, stabilization of particular conformational states^40,41^, or effects on enzymatic activity^42^.

To overcome these limitations, we created and validated a very large, fully randomized peptide library in a yeast display format, comprising ∼6.10 × 10^9^ unique peptides and ∼23% of total possible sequence space for peptides eight amino acids and shorter. Its diversity and unbiased design render this library suitable for obtaining peptide binders to a wide variety of targets. Here we present four selection campaigns demonstrating the broad utility of the library: two against IgG antibodies, one against an E3 ubiquitin ligase, and one against an essential membrane-bound bacterial enzyme involved in peptidoglycan synthesis. In each case we identified multiple high-affinity ligands, as well as motifs conserved among the highest affinity peptides, a result we anticipate will be generally true for a wide variety of protein targets.

## Results

### Library construction and validation

We sought to build a peptide library that can be used to screen a wide variety of protein targets for binding to a large proportion of all possible short peptides. By covering a large proportion of peptide sequence space, such a screen can identify not only individual peptide hits but also similar sequences or broader motifs common to peptide binders, providing insights into the determinants of a target’s binding interactions. To enable thorough sampling of sequence space in spite of practical constraints imposed by yeast transformation rates and culture volumes, we chose to limit the peptide length to a maximum of eight amino acids. Peptides of eight residues or fewer are common among biologically relevant protein-peptide interactions, such as antibody-epitope tag pairs (e.g., FLAG^43,44^, polyhistidine^45^, Strep-tag II^46^) and peptide G protein coupled receptor ligands (e.g., angiotensin II^47^, enkephalins^48^). Additionally, it is common for only a short fragment of a longer polypeptide to directly interact with a binding partner (e.g., the peptide apelin and its receptor APJ^49,50^). Thus, our library design allows us to sample a large fraction of the theoretical library diversity (i.e., sequence space) while still sampling biologically relevant peptide lengths.

Our library uses the Aga1-Aga2 display system^51^ to display peptides on the yeast cell surface, covalently anchored to the cell wall. We used eight degenerate NNK codons to confer diversity, resulting in the possibility of all 20 amino acids, as well as stop codons, at every variable position. A functional consequence of this design is that most clones (∼78%) will contain eight amino acids, but all shorter lengths are also sampled at lower frequencies. The lower theoretical diversity of shorter peptides compensates for their lower abundance, such that for most lengths effectively all possible sequences are represented in the library. The complete library expression construct encodes the Aga2 protein followed by a [G_4_S]_3_ linker, a hemagglutinin epitope tag (YPYDVPDYA) for monitoring expression, a single G_4_S linker, and finally the diversified peptide with a free carboxyl terminus (Fig. 1A).

**Figure 1.**
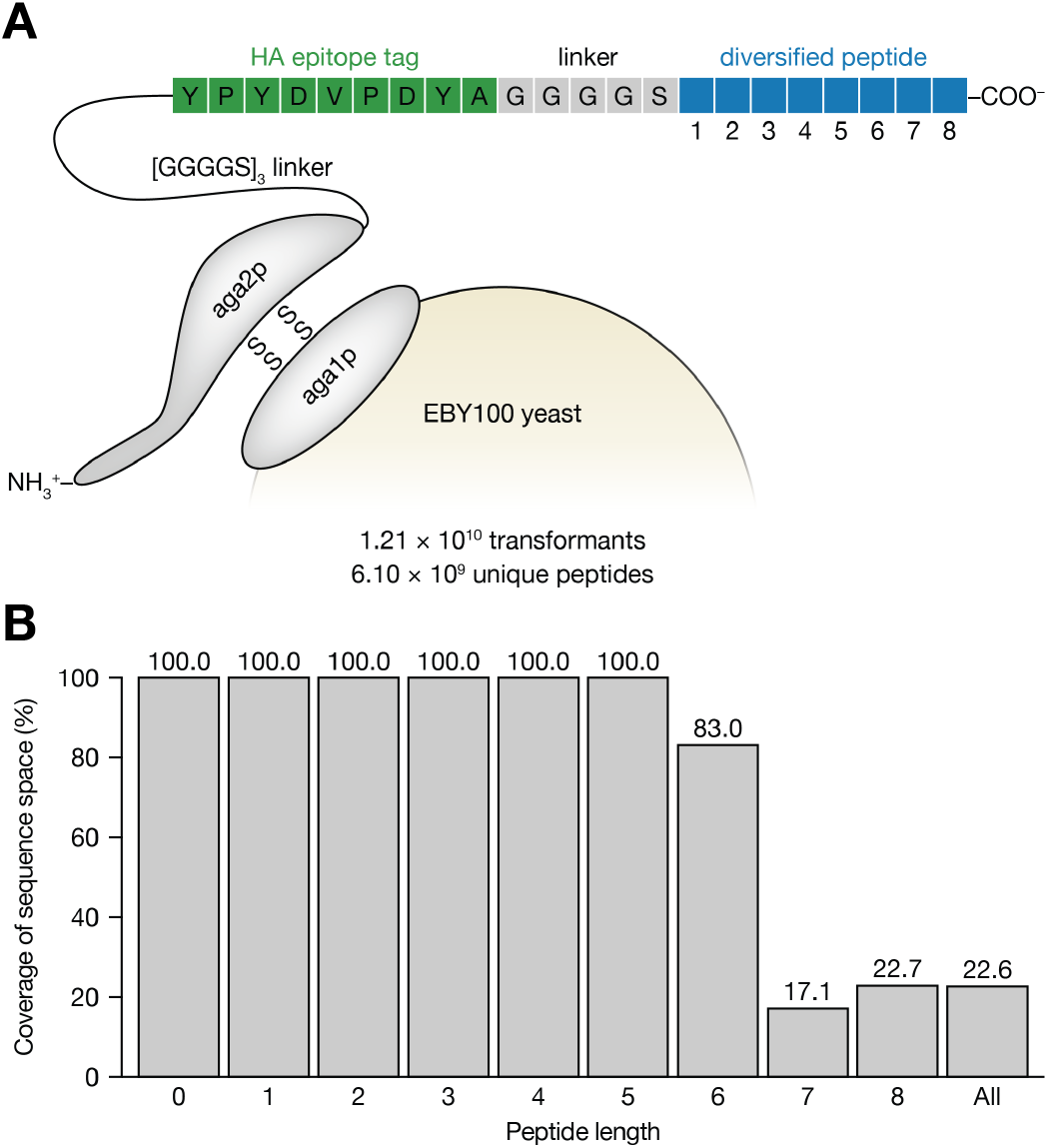
Design and assembly of a large, randomized peptide library. **A**, Schematic of yeast surface display expression construct. Diversified peptide region (blue) contains eight positions, each of which may encode any of the 20 common amino acids or a stop codon. **B**, Diversity analysis results from *in silico* library assembly simulation to assess library diversity and redundancy.

A yeast display library was assembled according to this design, resulting in ∼1.21 × 10^10^ transformants (see Methods). In many combinatorial libraries the theoretical diversity is much greater than achieved diversity, which implies that nearly all transformants are unique. In contrast, our library’s achieved diversity is similar to the theoretical diversity and many transformants are likely to be redundant. In order to account for this property and estimate the true diversity of the library (i.e., the number of unique peptide sequences), we performed an *in silico* simulation of the library assembly, using parameters associated with the library design (e.g., NNK codon frequencies). After accounting for sequence redundancies at the amino acid level, the simulated library’s diversity was ∼6.10 × 10^9^, corresponding to 22.6% of the theoretical diversity (i.e., all possible peptides eight amino acids and shorter). As a function of peptide length, this corresponds to 100% coverage of sequence space for peptides of length 0–5, 83.0% for six amino acid peptides, 17.1% for seven amino acid peptides, and 22.7% for eight amino acid peptides (Fig. 1B).

A sample of the final assembled library was used to prepare and sequence a Next Generation Sequencing (NGS) library to validate that the library matched its intended design. The observed peptide length distribution was very similar to that of the design (Fig. 2A). The amino acid usage distribution also closely matched the expected NNK codon distribution, both overall and at each of the eight variable positions (Fig. 2B and 2C). As a final validation measure, a sample of the library was stained with an anti-hemagglutinin antibody and analyzed by flow cytometry to assess peptide expression in the naïve library. We found that 67.0% of clones displayed the epitope tag above background (uninduced) levels (Fig. 2D).

**Figure 2.**
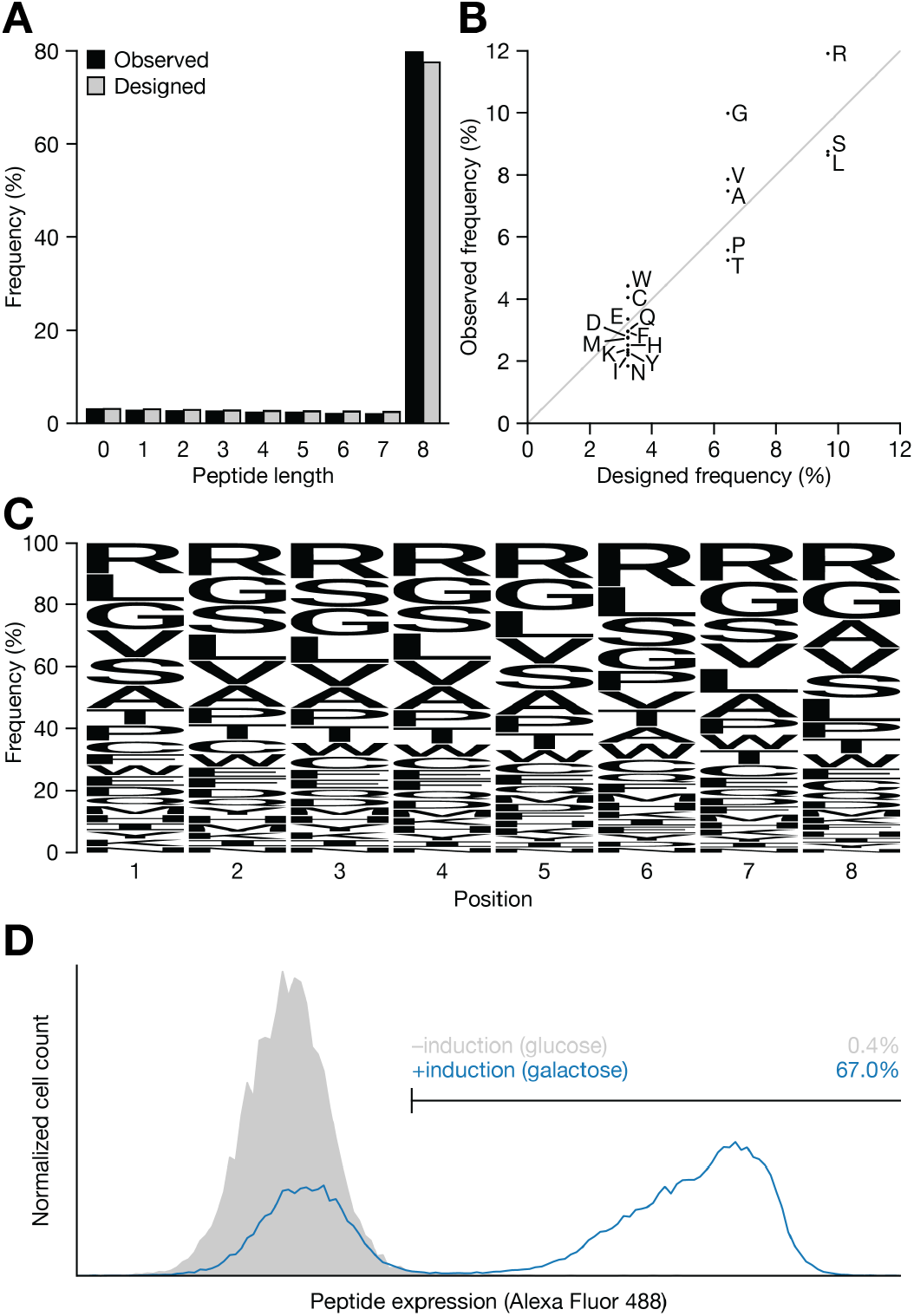
Experimental validation of assembled library. **A**, Histogram of peptide abundance in the library as a function of peptide length. **B**, Overall frequency of amino acid usage in the assembled library versus the library design (i.e., the NNK codon distribution). **C**, Per-position amino acid usage frequencies in the assembled library. **D**, Representative histogram of naïve yeast library expression as measured by binding to anti-hemagglutinin antibody.

### Rho1D4 selection campaign

To test the utility of the library, we performed a selection campaign against Rho1D4, an antibody discovered by mouse immunization with bovine rhodopsin that binds the C-terminal sequence ETSQVAPA with high affinity^52,53^. The goal of this campaign was to recover this canonical epitope or highly similar sequences. In this campaign, 10^10^ cells were sampled from the library and enriched for binding to biotinylated Rho1D4 antibody and counter-selected to prevent direct binding to streptavidin by Magnetic Activated Cell Sorting (MACS). Subsequently, the round 1 pool was further enriched by two rounds of Fluorescence Activated Cell Sorting (FACS) (Fig. 3A).

**Figure 3.**
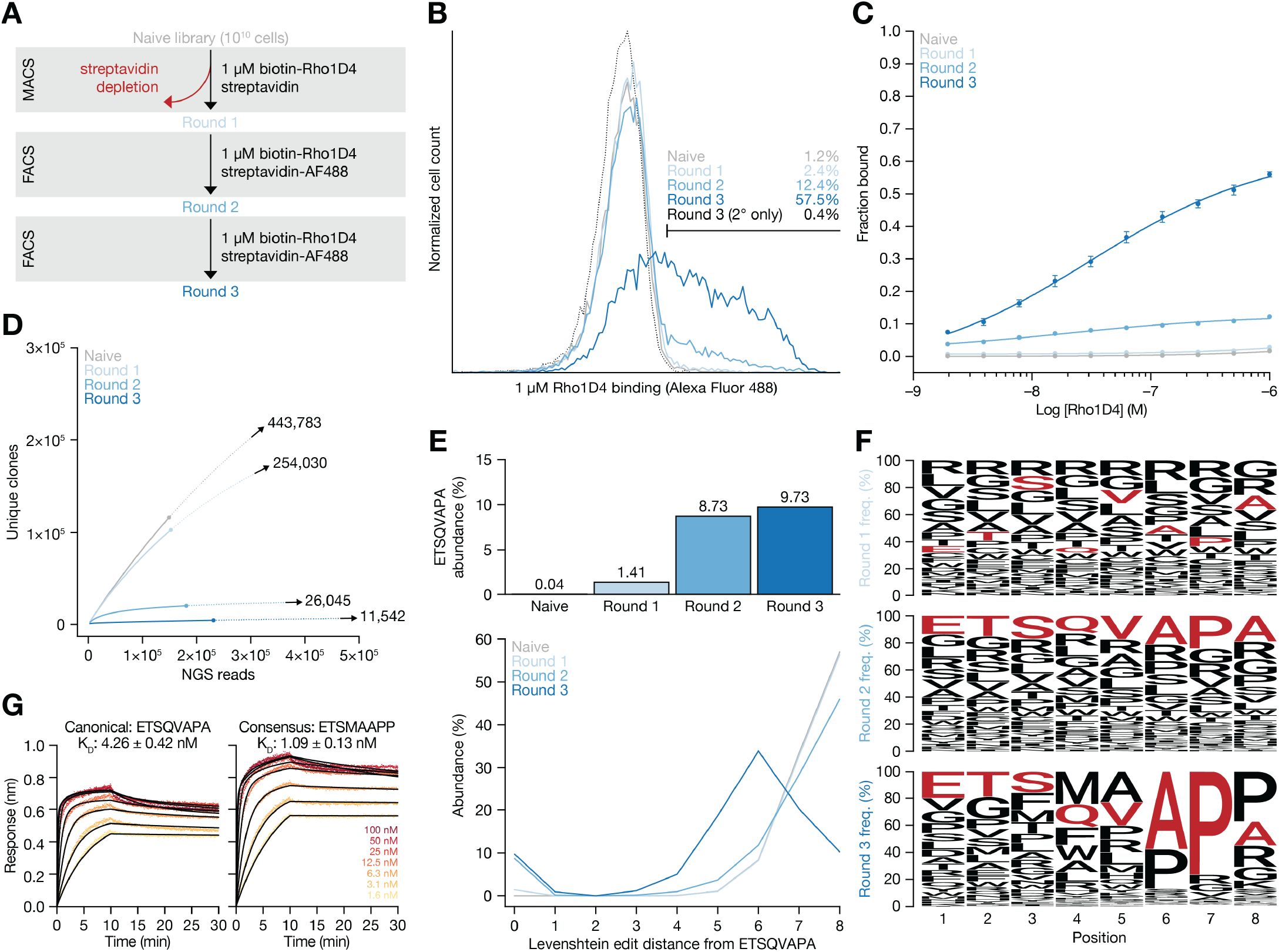
Selection campaign against the IgG antibody Rho1D4. **A**, Schematic of selection campaign. In MACS round, depletion and enrichment were performed sequentially. **B**, Representative histogram of per-round binding to 1 μM biotinylated Rho1D4-streptavidin Alexa Fluor 488 complex. **C**, Per-round Rho1D4 binding as assessed by flow cytometry in titration format. Data are presented as mean ± SEM from three independent technical replicates. **D**, Rarefaction curves (solid lines) for each round, with extrapolations (dotted lines) shown up to two-fold above the number of observed NGS reads. Chao1 diversity estimators for each round are presented numerically. **E**, The fraction of NGS reads representing the canonical Rho1D4 epitope (top). The fraction of NGS reads as a function of Levenshtein edit distance from the canonical Rho1D4 epitope (bottom). **F**, Sequence logos for each round. Residues corresponding to the canonical Rho1D4 epitope are highlighted in red. **G**, Representative biolayer interferometry sensorgrams for peptides corresponding to the canonical epitope (left) and the round 3 consensus sequence (right). Data are presented as mean kinetic fit K_D_ values ± SEM from three independent technical replicates.

At the end of the campaign, strongly enhanced binding to Rho1D4 was observed by flow cytometry, with 57.5% of round 3 cells binding above background in the presence of 1 μM Rho1D4, compared to 1.2% of cells in the naïve library (Fig. 3B). Intermediate rounds showed increasing binding throughout the campaign, across a range of concentrations from approximately 2 nM to 1 μM (Fig. 3C).

To assess the diversity at each round of the selection, we performed NGS sequencing and used these data to perform rarefaction-extrapolation analyses and calculate a Chao1 estimator, a non-parametric lower bound estimate of diversity^54,55^. By this method, it can be seen that diversity at the amino acid level was greatly reduced throughout the campaign, particularly during the two FACS rounds. Although the final round 3 pool shows strong interaction with Rho1D4, its relatively large Chao1 value of 11,542 suggests this pool could still be used for additional, more stringent selection rounds if desired (Fig. 3D).

Next, we quantified the frequency of the canonical Rho1D4 epitope, ETSQVAPA, in the NGS data for each selection round. After the first round of selection and in all subsequent rounds, the expected epitope sequence was the most abundant individual sequence, increasing from 0.04% in the naïve library to 9.73% in round 3 (Fig. 3E, top panel). The relatively high proportion in the naïve library was surprising, since the expected proportion based on NNK codon frequencies would be ∼0.000000009%. This discrepancy may be a result of bias in PCR amplification during NGS library preparation or cross-contamination with control yeast cells or a primer encoding this peptide. Since we expected to find not only the exact epitope sequence but potentially also closely related sequences, we assessed abundance in the NGS data as a function of edit distance from the canonical epitope. As expected, we also observed an increase in sequences with single-residue differences (0.00% in naïve, 0.91% in round 3), and a general shift toward lower edit distances throughout the campaign, indicating that the selection process enriched for similar sequences and depleted highly divergent sequences (Fig. 3E, bottom panel). In addition to ETSQVAPA being the most abundant individual peptide after selection, in round 2 each constituent amino acid of this peptide was the most common in its respective position (Fig. 3F, middle panel). In round 3 however, other amino acids appeared more frequently in positions 4, 5, and 8 (Fig. 3F, bottom panel) and many of the highest abundance individual peptides contained a C-terminal “APP” motif rather than the canonical “APA”.

To test whether these more frequent amino acid choices at positions 4, 5, and 8 confer higher affinity than the original immunogen sequence, we synthesized peptides corresponding to the canonical sequence as well as the round 3 consensus sequence (ETSMAAPP; 62.5% identity with the canonical epitope) and measured their binding to Rho1D4 using biolayer interferometry (BLI). The consensus sequence from the selection bound with a K_D_ value of 1.09 nM, more tightly than the original immunogen peptide, which had a K_D_ value of 4.26 nM (Fig. 3G). Interestingly, the consensus sequence did not appear as an individual peptide in any of the campaign NGS datasets, indicating that the ensemble data from a selection can be used to identify tightly interacting peptides without those peptides themselves being selected or even necessarily contained in the library. Additionally, the second-most abundant peptide in the round 3 NGS dataset (VGFWAPPK; 12.5% identity with the canonical epitope) was also synthesized and tested by BLI. It bound in the low micromolar range but was too weak to be reliably quantified.

### HPC4 selection campaign

Next, we screened the library against HPC4, an antibody known to bind a 12 amino acid epitope from human Protein C (EDQVDPRLIDGK), to test whether we could use the library to identify a shorter sequence that is nonetheless sufficient for high-affinity binding^56,57^. Such a sequence could be either a subsequence of the canonical epitope or an altogether novel sequence. The selection strategy was similar to that of the Rho1D4 campaign, with one round of MACS sorting and two rounds of FACS sorting (Fig. 4A). Binding was again strongly enriched throughout the campaign, with 95.6% of round 3 peptide-expressing cells binding HPC4 at 1 μM, compared to 1.4% in the naïve library (Fig. 4B). Strikingly, more than 50% of round 3 cells bound even at 2 nM, the lowest concentration tested (Fig. 4C).

**Figure 4.**
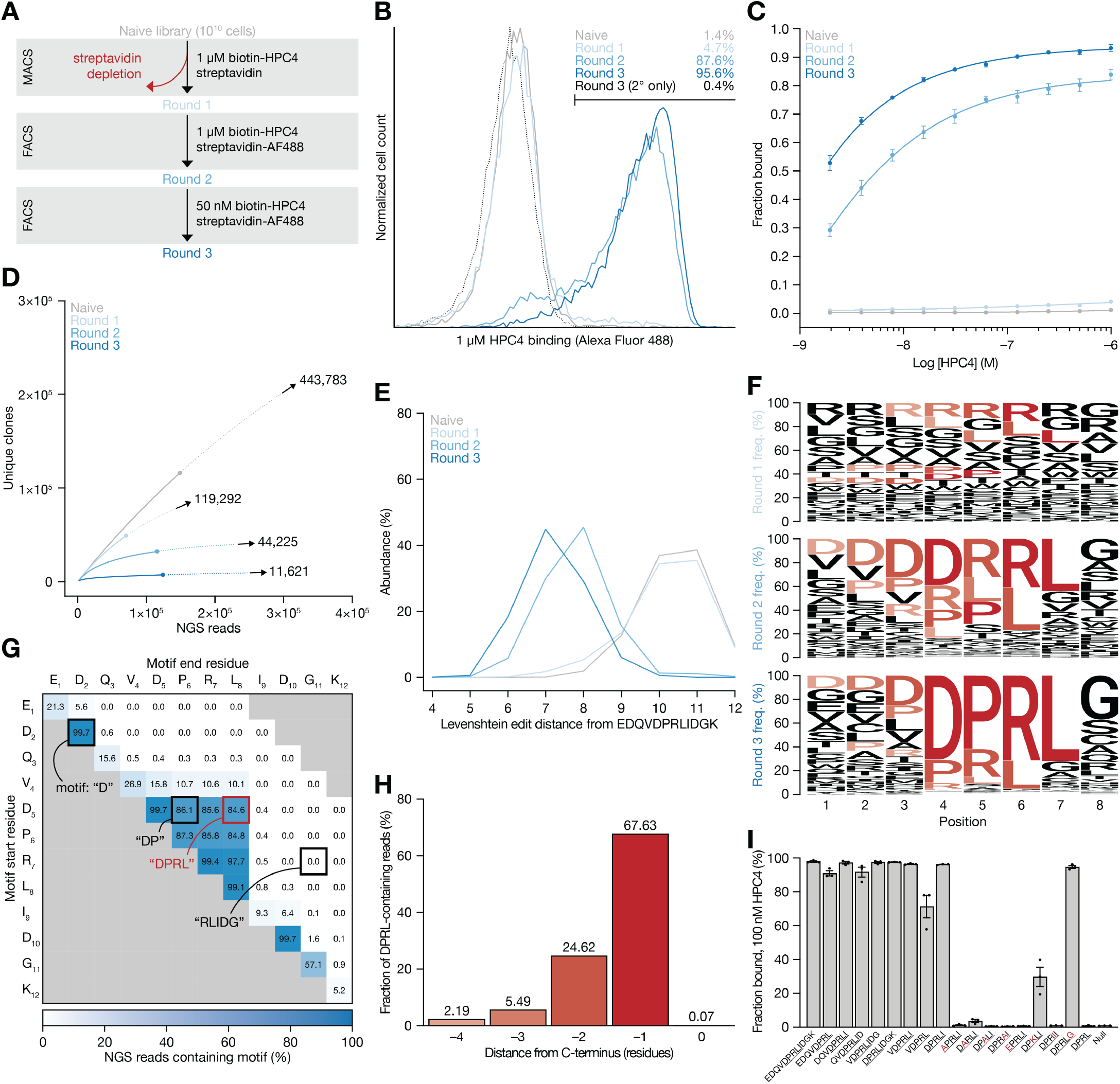
Selection campaign against the IgG antibody HPC4. **A**, Schematic of selection campaign. In MACS round, depletion and enrichment were performed sequentially. **B**, Representative histogram of per-round binding to 1 μM biotinylated HPC4-streptavidin Alexa Fluor 488 complex. **C**, Per-round HPC4 binding as assessed by flow cytometry in titration format. Data are presented as mean ± SEM from three independent technical replicates. **D**, Rarefaction curves (solid lines) for each round, with extrapolations (dotted lines) shown up to two-fold above the number of observed NGS reads. Chao1 diversity estimators for each round are presented numerically. **E**, The fraction of NGS reads as a function of Levenshtein edit distance from the canonical HPC4 epitope. **F**, Sequence logos for each round. The core “DPRL” motif is highlighted in shades of red, with shading corresponding to the register in which the motif appears. **G**, The fraction of NGS reads containing each possible subsequence of the canonical 12 amino acid HPC4 motif. Subsequence starting residue is designated on the y axis and ending residue on the x axis. Example subsequences are highlighted in black, and core “DPRL” motif is highlighted in red. **H**, Fraction of NGS reads containing “DPRL” motif as a function of the register in which the motif appears. **I**, Fraction of clonal yeast cells expressing variants of the HPC4 motif bound at 100 nM HPC4. Data are presented as mean ± SEM from three independent technical replicates.

As before, NGS libraries were prepared and sequenced at each round of selection. Rarefaction-extrapolation analysis was performed and used to tune selection pressure (i.e., target concentration and FACS gating strategy) to achieve a consistent rate of diversity reduction in each round (Fig. 4D). At the conclusion of the campaign, the round 3 Chao1 diversity estimate was 11,621 unique clones.

Although the canonical HPC4 epitope is 12 amino acids and therefore longer than any peptides contained within our library, we analyzed each round’s NGS dataset with respect to edit distance to the canonical epitope. In this case the minimum possible edit distance was four, representing an exact match to an eight amino acid subsequence of the longer canonical epitope. As with Rho1D4, we saw a leftward shift toward lower edit distances throughout the campaign, indicating depletion of highly divergent sequences and enrichment of sequences containing some similarity to the canonical epitope (Fig. 4E). However, unlike Rho1D4, we did not observe enrichment of exact matches to the canonical epitope, and instead saw the strongest enrichment at edit distances of 7 and 8. Sequence logos generated from each round’s NGS data indicated that these peaks were primarily attributable to a motif, “DPRL”, which corresponds to residues 5–8 of the canonical sequence (Fig. 4F). To more thoroughly examine this observation and investigate the possibility of less obvious enrichment of other subsequences, we calculated the proportion of round 3 NGS reads containing every possible subsequence of the canonical epitope. In this analysis, 84.6% of reads contained “DPRL”. Reads containing the slightly longer motif “VDPRL” comprised 10.1% of all reads, and no other highly enriched motifs were observed (Fig. 4G). Interestingly, the “DPRL” motif appeared in each of the five possible registers, most commonly one position from the C-terminus (i.e., xxxDPRLx) (Fig. 4H).

To probe the specificity of the peptide-HPC4 interaction, we tested several control peptides in a single-concentration flow cytometry assay (Fig. 4I). First, we tested binding of the full 12 amino acid epitope at 100 nM HPC4, and observed 97.9% of cells binding above background. Next, we tested each of the five possible eight amino acid subsequences of the canonical epitope (which notably all contain the “DPRL” motif), and again all exhibited nearly 100% binding (90.9– 97.6%). We then tested shorter truncations surrounding “DPRL”—96.5% of “VDPRLI” and 96.2% of “DPRLI” cells bound, and slightly reduced binding (71.2%) was observed with “VDPRL”, suggesting the residue immediately following this motif is more important for maintaining the binding interaction than the residue immediately preceding it. Next, single mutants of “DPRLI” were made, including mutating each of the four “DPRL” residues to alanine and three more subtle mutations (D→E, R→K, L→I). Each of these mutations substantially reduced binding, with all but the R→K mutation completely abolishing binding (<3.7% of cells bound). The R→K mutant retained 29.7% of cells bound. We then tested the mutant “DPRLG”, as glycine was the most commonly observed residue following the “DPRL” motif, and it too bound nearly maximally (94.7%). Finally, “DPRL” and the null peptide (i.e., the Aga2-based tether lacking a C-terminal peptide) were tested, and both showed no apparent binding (<1.0%). Together, these findings indicate that DPRL is indeed the core motif and is sensitive to any perturbations, but requires at least one additional amino acid to confer high-affinity binding. This additional residue may be N- or C-terminal, with C-terminal appearing to confer a greater enhancement to affinity and neither position appearing to have high amino acid specificity.

### KLHDC2 selection campaign

In the third selection campaign, we investigated whether the library can be used to identify recognition sequences (known as degrons) of an E3 ubiquitin ligase. Such an approach could be used to broadly profile ubiquitin ligase substrates and inform targeted protein degradation efforts. We chose as a target KLHDC2, an E3 ligase that has been shown to interact with the C-terminus of a premature translation product of the selenoprotein SelK (C-terminus: HLRGSPPPMAGG) and other substrates containing a C-terminal diglycine motif^58,59^. As many E3 ligase recognition motifs are short and therefore highly sampled in our library, we chose to substantially subsample the library in this selection, using only 2 × 10^7^ cells. This allowed us to proceed directly to FACS-based sorting, skipping the MACS debulking step performed in previous selection campaigns. We performed four rounds of FACS sorting, starting at 1 μM and lowering the concentration of target to 10 nM and 1 nM in rounds 3 and 4, respectively (Fig. 5A). A substantial improvement in binding was observed after even one round, which was improved further in subsequent rounds (Fig. 5B and 5C).

**Figure 5.**
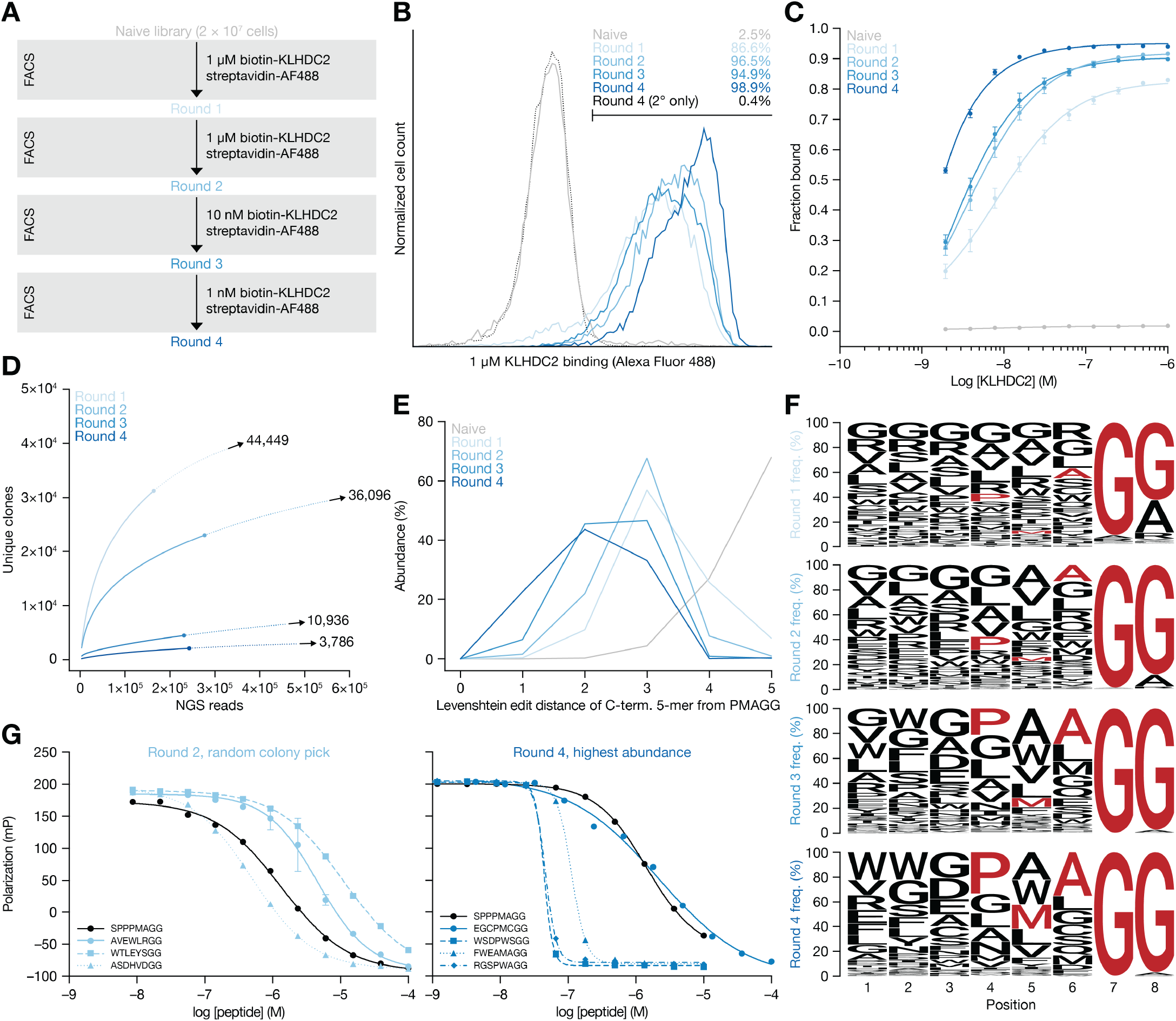
Selection campaign against the E3 ubiquitin ligase KLHDC2. **A**, Schematic of selection campaign. **B**, Representative histogram of perround binding to 1 μM biotinylated KLHDC2-streptavidin Alexa Fluor 488 complex. **C**, Per-round KLHDC2 binding as assessed by flow cytometry in titration format. Data are presented as mean ± SEM from three independent technical replicates. **D**, Rarefaction curves (solid lines) for each round, with extrapolations (dotted lines) shown up to two-fold above the number of observed NGS reads. Chao1 diversity estimators for each round are presented numerically. **E**, The fraction of NGS reads as a function of Levenshtein edit distance from the core C-terminal SelK recognition motif (PMAGG). **F**, Sequence logos for each round. Residues corresponding to the core SelK recognition motif are highlighted in red. **G**, Binding of selected peptides in fluorescence polarization assay. Clones chosen randomly from round 2 are shown on left (light blue), and highest abundance clones in the round 4 NGS dataset are shown on right (dark blue). SelK control 8-mer peptide is shown in black. Data are presented as mean ± SEM from three independent technical replicates.

As before, we performed NGS sequencing and rarefaction-extrapolation analysis at each round. This suggested a population of 3,786 unique clones in round 4, the vast majority of which we directly observed in the NGS data (Fig. 5D). In order to compare our results to the known recognition sequence from SelK, we truncated the peptide sequences in our NGS data to include only the five C-terminal residues and quantified their edit distances with respect to the previously shown^58^ core binding sequence of SelK (PMAGG-COO^-^). As before, we observe a general shift toward more similar sequences, with a peak at an edit distance of two (Fig. 5E). Strikingly, nearly all selected sequences in rounds 3 and 4 contain a diglycine motif, while the preceding “PMA” sequence is less conserved (Fig. 5F). This suggests that in agreement with previous studies, the diglycine motif is the primary driver of high-affinity interactions with KLHDC2, while the identities of the preceding residues may be less important. The most common residue other than glycine in positions 7 and 8 was alanine in position 8, occurring in 9.4%, 2.0%, and 2.5% of round 2, 3, and 4 NGS reads respectively. This observation agrees with previous findings that some sequences ending in GA or GS are also recognized by KLHDC2, in addition to the more common diglycine motif^59,60^.

To assess the success of the selection campaign, several selected clones were chosen, their corresponding peptides were synthesized, and their affinities were measured in competition fluorescence polarization assays. Three clones were chosen by random colony picking from round 2, and four clones with the highest abundance in the round 4 NGS dataset were selected. These clones demonstrated a range of affinities, with K_i_ values ranging from 36.8 nM to 9.4 μM, and notably one of three round 2 clones and three of four round 4 clones showed higher affinity than the SelK 8-mer (SPPP-MAGG) (Fig. 5G).

### RodA-PBP2 selection campaign

Our final selection was intended to test whether our library could be used for less tractable targets, for example membrane proteins that require detergent solubilization. Additionally, we sought to screen a target lacking any known peptide binding sites. Our test case for these questions was RodA-PBP2 from *E. coli*, an inner membrane-bound heterodimeric enzyme involved in the synthesis and maintenance of the peptidoglycan layer, and a complex for which there are no known peptide ligands. Previous work^61^ had identified sets of mutations that stabilize an active conformation of this enzyme, and one of these mutants (T52R, S54A, N55E; referred to simply as “RodA-PBP2” hereafter) was used throughout the selection campaign and in follow-up assays.

This campaign consisted of two rounds of MACS and two rounds of FACS, starting at 400 nM RodA-PBP2 and decreasing to 100 nM (Fig. 6A). Strong binding was observed by round 3 and further enhanced by round 4 (12.8% and 43.3% of cells bound at 400 nM, respectively) (Fig. 6B and 6C). Consistent reduction in diversity was observed throughout the campaign, with estimates of 14,180 and 8,392 unique clones in rounds 3 and 4 respectively (Fig. 6D). Examination of per-round sequence logos showed strong enrichment for tryptophan and other hydrophobic residues, raising the possibility that the campaign primarily enriched for nonspecific peptides that simply partition to the detergent micelle rather than *bona fide* RodA-PBP2 ligands (Fig. 6F). To assess this, the eight highest abundance peptides from round 4 were synthesized and tested for binding to RodA-PBP2 and a similar detergent-bound off-target, *E. coli* PBP1b (Fig. 6G, left panel). In this single-point BLI assay, two of the eight tested peptides exhibited binding specific to RodA-PBP2 but not the off-target protein, and follow-up assays showed that these peptides have K_D_ values of 1.14 μM and 0.76 μM. The two specifically-binding peptides, WAWEGTGP and FVFCGSGP share some chemical similarities, suggesting a common motif (i.e., Ω(A/V)ΩxGζGP) that could be investigated further to identify ligands with higher affinities or functional effects on RodA-PBP2. The binding properties of the eight tested peptides do not appear to be correlated with hydrophobicity, as calculated using the GRAVY index (Fig. 6G, right panel)^62^.

**Figure 6.**
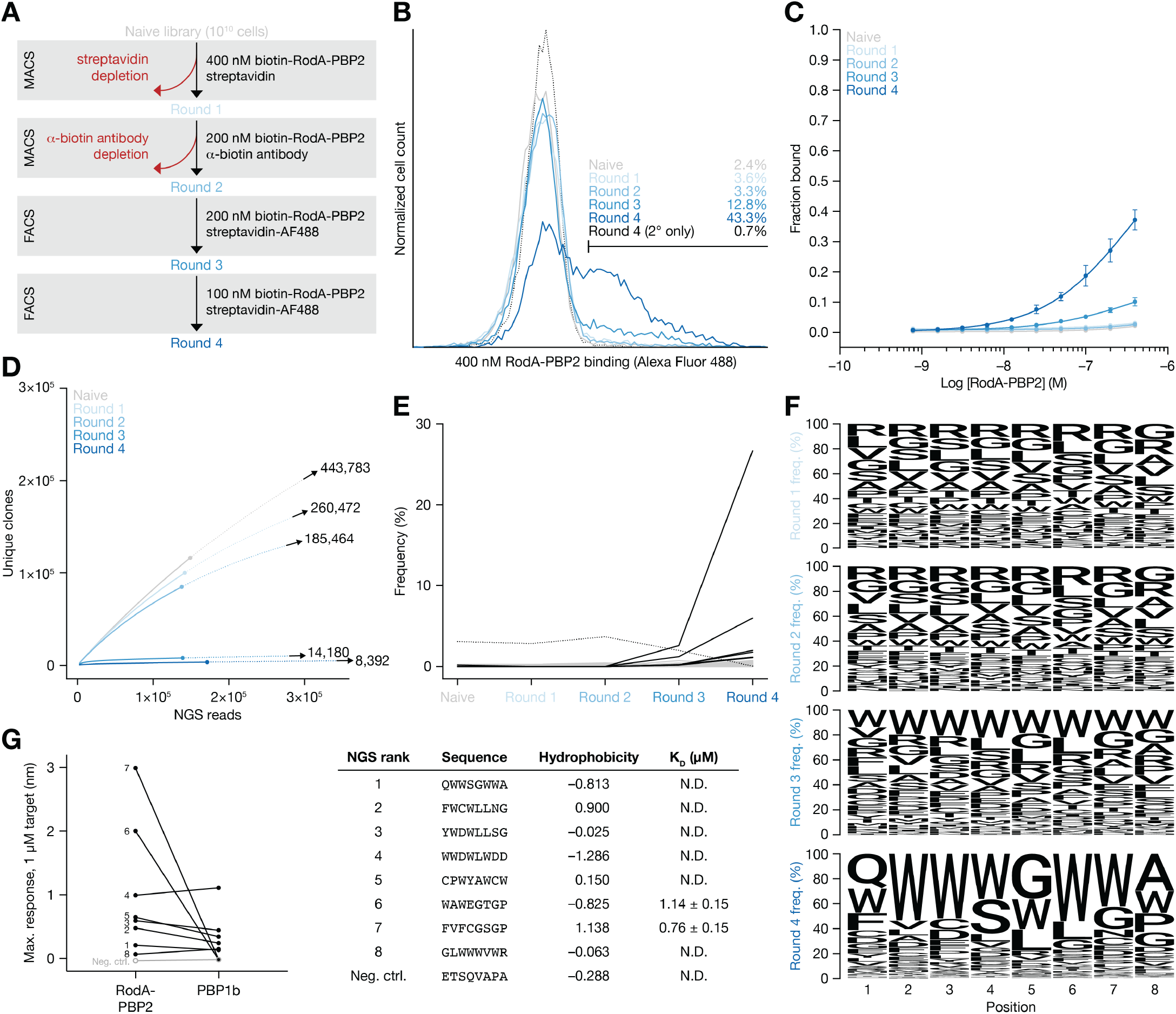
Selection campaign against the peptidoglycan synthase RodA-PBP2. **A**, Schematic of selection campaign. In MACS rounds, depletion and enrichment were performed sequentially. **B**, Representative histogram of per-round binding to 400 nM biotinylated RodA-PBP2-streptavidin Alexa Fluor 488 complex. **C**, Per-round RodA-PBP2 binding as assessed by flow cytometry in titration format. Data are presented as mean ± SEM from three independent technical replicates. **D**, Rarefaction curves (solid lines) for each round, with extrapolations (dotted lines) shown up to two-fold above the number of observed NGS reads. Chao1 diversity estimators for each round are presented numerically. **E**, Each peptide’s abundance in the NGS datasets for each selection round. The eight highest abundance sequences in round 4 are shown as solids black lines, the null peptide is shown as a dotted black line, and all other sequences are shown in gray. **F**, Sequence logos for each round. **G**, Binding of selected peptides in biolayer interferometry (BLI) assay. Single-concentration BLI responses for the eight highest abundance hits and negative control peptide against RodA-PBP2 and an off-target, PBP1b (left). Table showing the eight highest abundance hit sequences, their calculated hydrophobicities (GRAVY index), and their K_D_ values. K_D_ values are presented as mean equilibrium fit K_D_ values ± SEM from four independent technical replicates.

## Discussion

Although protein-peptide interactions are critical in biology, currently available tools to study such interactions are limited in several ways. In particular, peptide libraries are often biased toward a particular interaction of interest (e.g., deep mutational scan libraries) or particular sequences of interest (e.g., scanning proteome libraries) rather than being fully randomized. Additionally, many peptide libraries undersample peptide sequence space and therefore lack the scale required to broadly characterize the peptide-binding properties of a given target. By creating a peptide library using all amino acids at every variable position and at a scale approaching full coverage of sequence space, we aim to enable screens that can begin to identify the “rules” of peptide binding for a variety of protein targets going forward.

In this work, we describe the design, construction, and validation of a large peptide library and demonstrate several examples of screens using this library that answer different questions about the peptide-binding properties of the given protein target. We have generated a library of ∼6.10 × 10^9^ unique peptides—including ∼100% of peptides of length 0–5 and ∼23% of all possible peptides up to length eight—in a yeast surface display format and extensively validated it for successful use in selection campaigns. Additionally, mutations are often functionally synonymous, so the proportion of functional chemical space represented by the library is likely higher than the strict proportion of chemical space at a sequence level. As such, the library can be used as a starting point for further exhaustive screening around an initial hit.

We first performed a selection against the antibody Rho1D4 and recovered the expected canonical epitope, as well as several additional binding sequences with only modest sequence similarity. Notably, we identified a consensus sequence that bound Rho1D4 with ∼3.9× higher affinity than the canonical epitope, and this consensus sequence did not itself appear in in the selection, indicating that the population-level properties of a selected cell pool can provide insights into the determinants of binding even with incomplete sampling of sequence space.

We then investigated whether this library can be used to identify a minimal binding sequence of the antibody HPC4, a commonly used research antibody that binds the 12 amino acid Protein C epitope tag. In this campaign, we identified a core binding motif, DPRL, which is itself necessary but insufficient for high-affinity binding, and must have at least one additional preceding or following residue to confer tight binding. The Rho1D4 antibody, by contrast, recognizes a longer peptide epitope but tolerates amino acid changes at multiple positions.

Then we tested whether this library can be used to profile degron sequences of an E3 ubiquitin ligase, KLHDC2, which is known to recognize C-terminal degrons containing a diglycine motif. Our selection confirmed the importance of this diglycine motif for high-affinity interaction with KLHDC2, and we were able to isolate peptides with a range of affinities to the target, spanning K_i_ values from 36.8 nM to 9.4 μM. Notably, several of these peptides had affinities much higher than the C-terminus of the canonical substrate SelK.

Lastly, we tested whether this library can be used to identify novel peptide binders to a membrane protein, using as our target the *E. coli* RodA-PBP2 complex. In this campaign we were successful in discovering two novel peptide ligands, which bind to the RodA-PBP2 complex at ∼1 μM K_D_ and exhibit no binding to a similar bacterial penicillin binding protein, *E. coli* PBP1b.

These four selection vignettes represent examples of biological questions that can be investigated using our library and selection strategy, but there are countless additional questions for which this library could provide insight. For example, many antibodies in both research and clinical use interact with short linear epitopes, and this library could be used in a manner similar to the Rho1D4 and HPC4 selections to profile their sequence specificity and identify the core interacting residues. Likewise, other protein families such as E3 ubiquitin ligases and PDZ domains often interact with C-terminal peptide fragments and could be readily profiled using the methods described here^63^. In future studies, we plan to use this library to investigate the ligand space of various G protein coupled receptors. This work will aim to identify peptide ligands with unusual pharmacology or sequences highly divergent from known ligands, and could potentially serve as a starting point for deorphanization of receptors for which there are no known ligands.

As our yeast display library is amenable to FACS sorting, it could be adapted to answer more complex questions. For example, the use of labeled secondary reagents could enable multicolor FACS sorting schemes to identify peptides that bind at specific sites on the target, induce specific target conformations, or promote association with additional binding partners. Library-on-library screening approaches could also be taken to identify clones with nuanced properties^64^. In order to facilitate broad use of this library by external research groups, we have made it publicly available and free of charge for nonprofit use—it is available from a commercial vendor (Kerafast, Cat#EF0023-FP) or via Material Transfer Agreement.

While our library contains a large proportion of short linear peptide sequence space, its design necessarily precludes the discovery of certain classes of biologically important peptides, for example peptides containing non-canonical amino acids, peptides that contain a free N-terminus, cyclic peptides (other than those cyclized via disulfide bonds), and peptides requiring more than eight consecutive amino acids to facilitate their binding interactions. While these chemical classes are clearly important in both biology and medicine, they remain outside the scope of this platform, and warrant further research into new libraries and screening methods.

This platform is a new tool for profiling protein-peptide interactions across diverse protein targets, and features both near-comprehensive coverage of sequence space and flexibility to a wide range of biological questions. By making this resource available to the research community with minimal restrictions, we aim to enable future studies that will shed light on peptide binding mechanisms and facilitate tool compound and therapeutic discovery.

## Acknowledgements

We thank Dr. Timothy Springer for providing Rho1D4 and HPC4 hybridoma sequencing data. We thank Dr. Kelly Arnett at the Center for Macro-molecular Interactions in the Department of Biological Chemistry and Molecular Pharmacology at Harvard Medical School for assistance with biolayer interferometry experiments. We thank Dr. Meredith Skiba for critical reading of the manuscript. R.P.N. is a member of the excellence cluster ImmunoSensation2 funded by the Deutsche Forschungsgemeinschaft (DFG) under Germany’s Excellence Strategy – EXC2151–390873048. This work was funded by National Institutes of Health TR01 grant 1R01CA260415 to A.C.K., T32 training grant 5T32GM007226-46 to J.D.H., and Fujifilm Graduate Fellowship to J.D.H. This preprint was typeset with the bioRxiv word template by @Chrelli: www.github.com/chrelli/bioRxiv-word-template

## Author contributions

J.D.H. and A.C.K. conceived of the project. A.C.K., E.S.F., and R.P.N. supervised the project. J.D.H. built and validated the peptide library. J.D.H., I.S., M.L., H.Y., and Z.Z. prepared protein reagents. J.D.H., I.S., and Z.Z. performed MACS and FACS selections. J.D.H. performed flow cytometry assays. J.D.H. performed NGS sequencing and analysis. J.D.H. performed biolayer interferometry experiments. M.L. performed fluorescence polarization assays. J.D.H. wrote the manuscript with input from all authors.

## Data availability

NGS sequencing data have been deposited to GitHub (https://github.com/kruselab/peptide-library) and are publicly available as of the date of publication at https://doi.org/10.5281/zenodo.14947976.

## Code availability

All original code has been deposited to GitHub (https://github.com/kruse-lab/peptide-library) and is publicly available as of the date of publication at https://doi.org/10.5281/zenodo.14947976.

## Materials availability

All unique reagents generated in this study are available from the lead contact without restriction, other than the peptide library itself, which can be ordered directly (Kerafast, Cat#EF0023-FP) or obtained from the lead contact with a completed Materials Transfer Agreement.

## Materials and Methods

### Peptide library construction

A DNA library encoding diversified peptide sequences and flanking sequences for homologous recombination was constructed by assembly PCR. Overall length of insert and overhang lengths were optimized to obtain maximal transformation efficiency. A total of eight primers were used, including a single degenerate primer incorporating eight NNK codons that confer library diversity (Table S1). 50 μM assembly PCR reactions were prepared using 10 μM outer primers (Insert F1 and R1), 0.1 μM internal primers, and Q5 Hot Start High-Fidelity polymerase (NEB). Band corresponding to the full-length insert was gel extracted using QIAquick Gel Extraction Kit (Qiagen) and subsequently amplified in 50 μL reactions using 0.5 μM outer primers and Q5 Hot Start High-Fidelity polymerase. In both PCRs, template concentrations were chosen to ensure >100-fold excess with respect to the maximum theoretical library diversity. PCR product was purified using QIAquick PCR Purification kit (Qiagen) and subsequently ethanol precipitated prior to yeast transformation.

Expression plasmid was generated by modifying pYD1^65^ to add an HA tag (YPYDVPDYA) N-terminal of the displayed peptide and remove all other tags. This plasmid was linearized for yeast transformation by PCR using KOD Xtreme Hot Start polymerase (Millipore Sigma) and 0.3 μM primers (Vector F and R, Supplementary Table 1). PCR product was purified using QIAquick PCR Purification kit (Qiagen) and subsequently ethanol precipitated prior to yeast transformation.

To prepare electrocompetent yeast cells, a YPAD culture was inoculated from a single EBY100 colony and grown shaking at 30 °C. For each electroporation plate, a 1.1 L YPAD culture was inoculated at an initial OD600 of 0.3 and grown to mid-log phase (OD_600_ between 1.5–3). Culture was distributed into 50 mL conical tubes, pelleted by centrifugation at 2,000 *x g* for 5 min, then each tube was resuspended in 25 mL 100 mM lithium acetate, 10 mM DTT and incubated shaking at 30 °C for 10 min. Cells were pelleted at 2,000 *x g* for 5 min, then washed with 25 mL ice-cold water per tube. Supernatant was discarded and cells were resuspended in a total of 3.96 mL DNA mixture (105.6 μg linearized backbone and 211.2 μg insert dissolved in ice-cold water). 100 μL cell-DNA mixture was dispensed into each well of a 96 well electroporation plate with 2 mm gaps (BTX) and pulsed for 15 ms at 500 V using an ECM 830 Square Wave Electroporator (BTX). Cells were recovered in 480 mL YPAD for 60 min, then washed with selective medium lacking tryptophan, serially diluted, and plated on dropout medium to estimate transformation efficiency. After 48 hours, a quantity of cells corresponding to ten times the number of transformants was passaged into YGLC medium (0.38% -trp dropout media supplement (US Biological), 0.67 yeast nitrogen base, 2% glucose, 1.04% sodium citrate, 0.44% citric acid monohydrate, 1% (v/v) penicillin/streptomycin, pH 4.5) and grown for 24 hours at 30 °C. After 24 hours, the same quantity of cells was centrifuged at 3,500 *x g* for 5 min, resuspended in 90% YGLC/10% DMSO, and slow-frozen in cryovials.

### Diversity and redundancy simulations

To account for the presence of redundant peptides and more accurately estimate library diversity (i.e., the number of unique peptides), library assembly simulations were performed to estimate the number of unique transformants that would likely be obtained for a given number of total transformants. Sequences were generated such that the amino acid usage and peptide length distributions matched the theoretical library design (Fig. 2A and 2B) and a total of 1.21 × 10^10^ sequences were generated, corresponding to the number of transformants experimentally obtained. Redundant sequences were removed and unique sequences were counted, resulting in an estimate of library diversity.

### Protein reagent preparation

Rho1D4 antibody was produced from murine Rho1D4 hybridoma cells, which were continuously cultured in a CELLine disposable bioreactor (Corning), with approximately 40 mL of culture being removed from the chamber every 3-4 days. Culture was centrifuged at 250 *x g* for 5 min and supernatant was flash-frozen and stored at -20 °C for later purification. Supernatant was then thawed, mixed in 1:1 ratio with HBS pH 7.5 buffer, loaded onto 1D4 peptide resin (TETSQVAPA) by gravity flow, washed with HBS pH 7.5 buffer, eluted with 5 mM NaOH, and immediately neutralized with 1 M HEPES pH 7.5 buffer. Eluate was dialyzed against HBS pH 7.5 using 10 kDa Slide-a-Lyzer G3 cassettes (ThermoFisher). Rho1D4 antibody was biotinylated using NHS-LC-biotin (ThermoFisher) at a ratio of 10:1 biotins:IgG, and degree of labeling was measured using Pierce Biotin Quantitation Kit (ThermoFisher). Biotinylated protein was flash-frozen and stored at -80 °C for later use.

HPC4 antibody was produced from murine HPC4 hybridoma cells, which were continuously cultured in a CELLine disposable bioreactor (Corning), with approximately 40 mL of culture being removed from the chamber every 3-4 days. Culture was centrifuged at 250 *x g* for 5 min and supernatant was flash-frozen and stored at -20 °C for later purification. Supernatant was then thawed, mixed in 1:1 ratio with HBS pH 7.5 buffer and CaCl_2_ was added to a final concentration of 2 mM. Sample was loaded onto Protein C peptide resin (EDQVDPRLIDGK) by gravity flow, washed with HBS pH 7.5 buffer supplemented with 2 mM CaCl_2_, eluted with 100 mM sodium citrate pH 3, and immediately neutralized with 1 M HEPES pH 8 buffer. Eluate was dialyzed against HBS pH 7.5 using 10 kDa Slide-a-Lyzer G3 cassettes (ThermoFisher). HPC4 antibody was biotinylated using NHS-LC-biotin (ThermoFisher) at a ratio of 3:1 biotins:IgG, and degree of labeling was measured using Pierce Biotin Quantitation Kit (ThermoFisher). Biotinylated protein was flash-frozen and stored at -80 °C for later use.

Human KLHDC2 (kelch domain, amino acids 1–362) was cloned into a pAC-derived vector and recombinant protein was expressed as a N-terminal Strep tag fusion in *Trichoplusia ni* High-Five insect cells using the baculovirus expression system (Invitrogen). Cells were pelleted by centrifugation and resuspended in lysis buffer containing 50 mM Tris pH 8.0, 200 mM NaCl, 0.1% Triton X-100, 1 mM Tris-(2-carboxyethyl) phosphine (TCEP), 1 mM phenylmethylsulfonyl fluoride (PMSF), and 1× of each of the protease inhibitors including Bestatin (10 μM), Leupeptin (10 μM), Aprotinin (2 μg/ml), E-64 (2 μM), Pepstatin (1 μM), and Phosphoramidone (1 μM). The cell suspension was sonicated followed by ultracentrifugation to clarify lysate. The soluble fraction was collected and flowed over Strep-Tactin XT Superflow (IBA) affinity resin, washed with wash buffer containing 50 mM Tris pH 8.0, 200 mM NaCl, 1 mM TCEP, and eluted with buffer containing 50 mM Tris pH 8.0, 200 mM NaCl, 1 mM TCEP, and 50 mM biotin. The protein was then biotinylated overnight in the presence of 10 mM MgCl_2_, 20 mM ATP, and 145 μg BirA enzyme per 5 mL reaction (biotin was already present in the elution buffer). The eluate was collected, concentrated, and injected onto Superdex 200 10/300 size-exclusion chromatography column (Cytiva) in a buffer containing 50 mM HEPES pH 7.4, 200 mM NaCl, and 1 mM TCEP. Elution fractions were pooled, concentrated, flash-frozen, and stored at –80 °C for later use.

*E. coli* RodA-PBP2 complex was purified as described previously^61^. Briefly, the plasmid encoding the active state-stabilized variant of the *Ec-* RodA-PBP2 fusion (pMG63: ColA-*T7::*His6-SUMO-Flag-3C-*Ec*RodA-[GGGS]_3_-*Ec*PBP2(T52R-S54A-N55E-V143T-A147S) was transformed into *E. coli* C43 (DE3) cells with SUMO tag-specific Ulp1 protease under an arabinose inducible plasmid (pAM174) and grown in Terrific broth (TB) medium supplemented with 50 μg/mL kanamycin, 35 μg/mL chloramphenicol, 2 mM MgCl_2_, 0.1% (w/v) glucose, and 0.4% (v/v) glycerol to an OD_600_ between 2–3 at 37 °C with vigorous shaking. Protein expression was induced with 1 mM IPTG and 0.1% arabinose at 18 °C. After 12–16 hours of induction, cultures were harvested by centrifugation and resuspended in 150 mL of lysis buffer per liter of culture (50 mM HEPES pH 7.5, 150 mM NaCl and 20 mM MgCl_2_, benzonase nuclease at 1:100,000 dilution, EDTA-free protease inhibitors). Cells were lysed with LM10 Microfluidizer (Microfluidics) and membrane fraction was isolated by centrifugation at 50,000 *x g* for 1 hour and solubilized in 150 mL of resuspension buffer per liter of culture (50 mM HEPES pH 7.5, 350 mM NaCl, 10% (v/v) glycerol, 1% (w/v) n-dodecyl-β-D-maltoside). Samples were stirred for 1–2 hours at 4 °C then centrifuged as before. The resulting supernatant was supplemented with 2 mM CaCl_2_ and loaded onto M1 anti-FLAG affinity resin equilibrated with wash I buffer (resuspension buffer supplemented with 2 mM CaCl_2_). The resin was subsequently washed with wash I buffer, followed by wash II buffer (20 mM HEPES pH 7.5, 350 mM NaCl, 5% (v/v) glycerol, 0.1% DDM, 2 mM CaCl_2_). Protein was eluted with 0.2 mg/mL FLAG peptide and 5 mM EGTA and concentrated to 1–5 mg/mL using a 100 kDa cutoff Amicon concentrator (Millipore Sigma). Protein was then injected onto a Superdex200 Increase size exclusion column (Cytiva) in a buffer containing 20 mM HEPES pH 7.5, 350 mM NaCl, and 0.1% (w/v) DDM. Elution fractions were pooled, concentrated, and biotinylated for 30 minutes at room temperature with EZ-Link NHS-LC-Biotin (ThermoFisher) added in 5-fold molar excess to the protein. Following biotinylation, excess biotin was removed through an additional pass on Superdex200 Increase column. Protein peak fractions were collected, concentrated, flash-frozen, and stored at -80 °C for further use. Enzymatic activity was assessed using the previously described polymerization assay^61^, and biotinylation efficiency was quantified with a gel-based streptavidin shift assay. Typically, this protocol yielded 80–90% biotinylated protein that retained polymerization activity.

### Cell sorting

For initial MACS cell sorting, the peptide library was subsampled at a depth of 10^10^ cells and grown in -trp +glu medium shaking at 30 °C. For RodA-PBP2 MACS 2 sorting, 4.1 × 10^8^ cells were used, corresponding to ten times the number of cells eluted in the previous MACS round. After 24–48 hours, the same number of cells was passaged into -trp +gal inducing medium and grown shaking at 25 °C for 48 hours. MACS sorting was performed according to manufacturer’s protocol (Miltenyi). Briefly, cells were pelleted at 3,500 *x g* for 5 min and resuspended in 10 mL selection buffer (protein storage buffer supplemented with 0.1% (w/v) BSA and 10 mM maltose). Cells were pelleted as before, resuspended in 4.5 mL selection buffer, and 500 μL of magnetic beads (streptavidin for all MACS rounds except RodA-PBP2 MACS 2, which used anti-biotin; Miltenyi) were added to the solution. Cell-bead mixture was incubated for 30 min at 4 °C, washed once more with 5 mL of selection buffer, then applied to an LD column on magnetic rack (Miltenyi) pre-equilibrated with selection buffer. Flowthrough was collected, pelleted as before, and resuspended in 5 mL selection buffer containing protein target (1 μM for Rho1D4 and HPC4, 400 nM and 200 nM for RodA-PBP2 MACS 1 and 2, respectively). Solution was incubated at 4 °C for 1 hour, then washed once and resuspended in selection buffer supplemented with 500 μL magnetic beads. Solution was incubated for 20 min, washed twice in 5 mL selection buffer, then applied to LS column on magnetic rack (Miltenyi) pre-equilibrated with selection buffer. LS column was washed with 8 mL selection buffer, then removed from magnetic rack and bound cells were eluted with provided plunger in 5 mL selection buffer. Cell count was estimated using flow cytometer, then eluate was pelleted and resuspended in 3 mL -trp +glu medium and recovered for 24 hours at 30 °C.

For FACS cell sorting, a number of cells corresponding to at least ten times the number collected in the previous selection round (or 2 × 10^7^ cells for KLHDC2 FACS 1 round) were grown in -trp +glu medium for 24–48 hours shaking at 30 °C. The same number of cells were passaged into -trp +gal inducing medium and grown for 48 hours shaking at 25 °C. The same number of cells were then pelleted at 3,500 *x g* for 5 min, resuspended in selection buffer (protein storage buffer supplemented with 0.1% (w/v) BSA and 10 mM maltose), pelleted once more, then resuspended at a density of 10^7^ cells/mL in selection buffer supplemented with protein target, Alexa Fluor 488 streptavidin (BioLegend) at a stoichiometry of 0.225× with respect to target, and 10 μg/mL antiHA-AF647 antibody. Target protein concentrations were generally chosen such that ∼20% of previous-round cells showed binding at the chosen concentration in bulk flow cytometry assays. Cells were sorted on Sony SH800 FACS using 100 μm chip and recovered in 3 mL -trp +glu medium shaking at 30 °C for 24–48 hours.

Following each selection round, at least ten times the number of cells collected were passaged into YGLC medium, grown for 24–48 hours at 30 °C, pelleted at 3,500 *x g* for 5 min, resuspended in YGLC supplemented with 10% DMSO, and slow frozen at -80 °C.

### Yeast control generation

Control plasmids were generated from library plasmid linearized with KOD Xtreme Hot Start polymerase (Millipore Sigma) and single oligos encoding peptide sequence surrounded by 25 nucleotide flanking sequences. Assembly was performed with NEBuilder HiFi DNA Assembly Master Mix (NEB). EBY100 yeast cells were transformed as described previously^66^.

### Flow cytometry

Stocks of yeast pools (at a cell count of at least ten times the number of cells collected in the previous selection round) or individual clones were thawed and cultured in -trp +glu medium for 24–48 hours. Cells were then induced in -trp +gal medium for 48 hours, stained in selection buffer supplemented with 10 μg/mL Alex Fluor 647-labeled anti-HA antibody, protein target, and Alexa Fluor 488-labeled streptavidin (BioLegend) at a stoichiometry of 0.225× with respect to target for 60 minutes. Cells were washed twice with selection buffer, then analyzed on Beckman Coulter CytoFLEX flow cytometer. Flow cytometry data were analyzed using custom scripts utilizing the flowCore and sp packages. Fraction bound metric represents the number of peptide-expressing cells binding above unstained cells divided by the total number of peptide-expressing cells.

### Next generation sequencing

The naïve library and each selected pool following MACS or FACS selection was cultured in -trp +glu for 24–48 hours, then 1 mL of lag phase culture (OD_600_ of approximately 6, corresponding to ∼10^8^ cells) was collected and plasmid DNA was extracted using Yeast Miniprep II kit (Zymo). Peptide region was amplified by PCR using Q5 Hot Start High-Fidelity polymerase (NEB; primers Insert F1 and R1, Supplementary Table 1). PCR product was purified using QIAquick PCR Purification kit (Qiagen), concentration-normalized using Qubit dsDNA HS kit (Invitrogen), and sequenced with Amplicon-EZ service (GeneWiz).

Paired-end reads were merged using NGmerge, then merged reads were processed using custom code. Briefly, reads were filtered for those matching the primers used for library generation and whose translated sequences matched the library design. Amino acid residues flanking the diversified peptide region were removed, and the peptide sequences alone were used for subsequent analyses.

Accumulation-rarefaction analysis was performed using the immunarch package. In particular, rarefaction-extrapolation curves were generated with the repDiversity() function using .method = “raref” and extrapolations truncated at twice the observed number of sequences, and Chao1 estimators were calculated using the repDiversity() function with parameter .method = “chao1”. Edit distance analysis was performed using the stringdist package with parameter method = “lv”. To generate sequence logos, first gaps were inserted such that all sequences were aligned at the C-terminal residue, then logos were generated using the geom_logo() function in the ggseqlogo package.

### Biolayer interferometry

Biotinylated peptides (GenScript; biotinylated at N-terminus and containing an N-terminal G_4_S linker) in protein storage buffer supplemented with 10% DMSO were immobilized on streptavidin biosensors (Sartorius), then washed in protein storage buffer. Sensors were then dipped into a 96- or 384-well black, flat-bottom plate (Grenier) containing protein target in protein storage buffer. All assays were performed at room temperature. For Rho1D4, kinetic fits were obtained in Octet Analysis Studio software and plotted in GraphPad Prism version 10. Reported K_D_ values are the mean ± SEM of three independent replicates. For RodA-PBP2, equilibrium responses were extracted and fit to a one site binding model in GraphPad Prism version 10. Reported K_D_ values are the mean ± SEM of four independent replicates.

### Fluorescence polarization assays

Competitor peptides (GenScript) were dispensed into a 384-well microplate (Corning, 4514) using a D300 Digital dispenser (HP) normalized to 1% DMSO into a final assay volume of 15 μL. Assay buffer contained 10 nM SelK 8mer (SPPPMAGG) labeled with FITC-Ahx (GenScript), 142 nM KLHDC2, 50 mM HEPES pH 7.5, 200 mM NaCl, 0.1% Pluronic F68, and 1 mM TCEP. Fluorescence polarization was measured using a PHERAstar FS microplate reader (BMG Labtech) for 1 hour, 42 secs in 30 cycles. All assays were performed at room temperature. Data from three independent replicates were plotted and IC_50_ values were estimated using a four-parameter inhibition model in GraphPad Prism version 10. The Cheng-Prusoff equation was applied to convert IC_50_ values into their corresponding K_i_ values.

