## Supplementary Information for "Decoding protein-peptide interactions using a large, target-agnostic yeast surface display library"

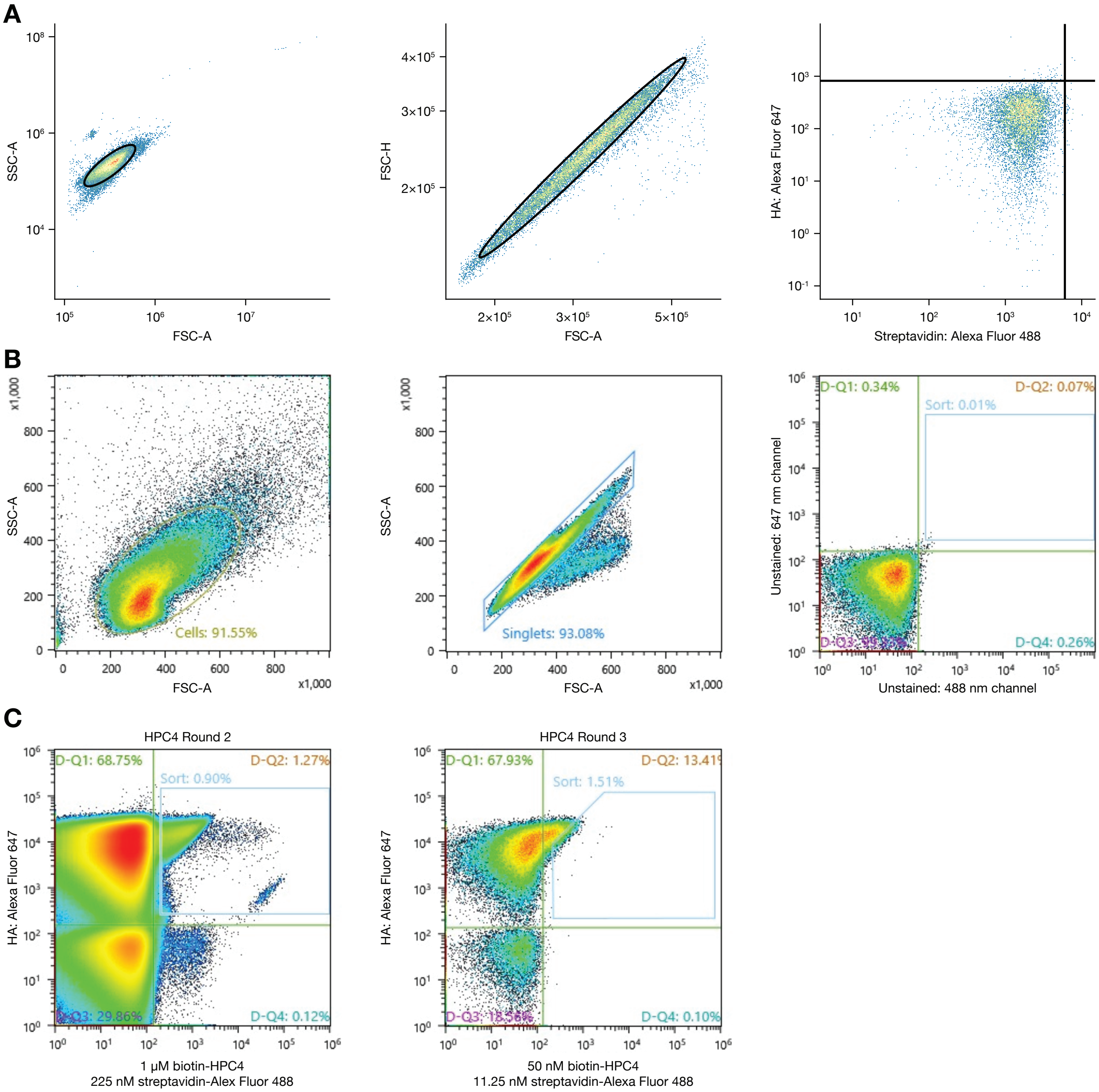


**Figure S1.** Representative flow cytometry gating strategy. **A**, Representative plots illustrating gating strategy for flow cytometry experiments (gates indicated by thick black lines). Data are shown for uninduced yeast cells that lack any peptide displayed on the cell surface. **B**, Representative plots illustrating gating strategy for FACS sorting experiments. Data are shown for uninduced yeast cells that lack any peptide displayed on the cell surface. **C**, Representative plots illustrating FACS sorting strategy. A rectangular gating strategy (as seen in the left panel) was used in first FACS round with the goal of collecting nearly all binding clones, and a more stringent pentagon gating strategy (as seen in the right panel) was used in subsequent rounds with the goal of collecting only clones with the highest binding relative to their expression level.

**A**

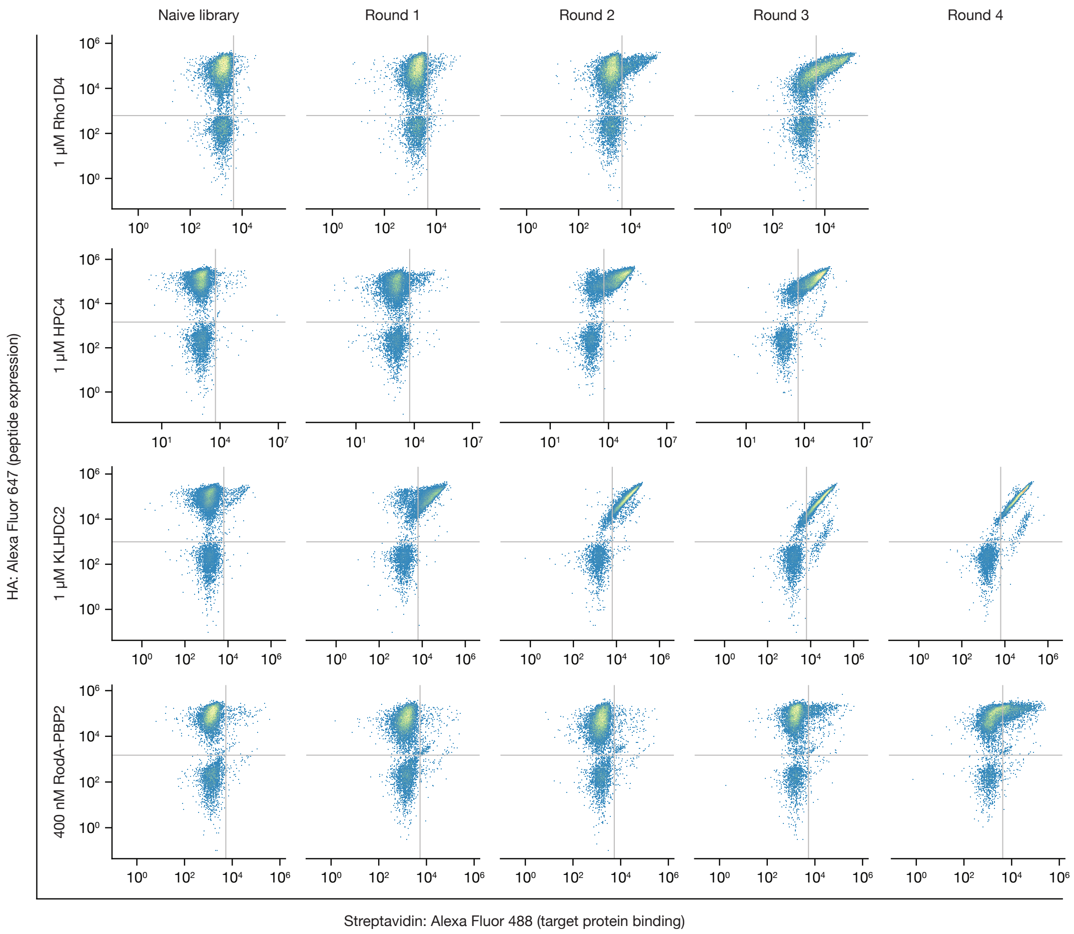


**Figure S2.** Representative flow cytometry of peptide selection rounds. **A**, Flow cytometry plots illustrating peptide expression and target protein binding throughout each selection campaign.

**Table S1.** Primers used in this study.

| Primer | Sequence |
|---|---|
| Insert F1 | CAAGGAGTTTTTGAATATTACAAATCAGTAACGTTTGTCAGTAATTGCGGTTCTCACCCCTCAACGACTAGCAAAGGCAGC |
| Insert F2 | ACACAGTATGTTTTTAAGCTTCTGCAGGCTAGTGGTGGTGGTGGTTCTGGTGGTGGTGGTTCTGGTGGTGGTGGTTCTTATCCGTATGA |
| Insert F3 | GGATTATGCGGGTGGTGGTGGTTCTNNKNNKNNKNNKNNKNNKNNKNNKTGAGTTTAAACCCGCTGATCTGATAACAACAGTG |
| Insert F4 | ACTTTGTTCCCACTGTACTTTTAGCTCGTACAAAATACAATATACTTTTCATTTCTCCGTAAACAACATGTTTTCCCATGTAATATCC |
| Insert R1 | AAAGTATGTGTAAAGTTGGTAACGGAACGAAAAATAGAAAAGGATATTACATGGGAAAACATGTTGTTTACGGAGAAATG |
| Insert R2 | GTATTTTGTACGAGCTAAAAGTACAGTGGGAACAAAGTCGATTTTGTTACATCTACACTGTTGTTATCAGATCAGCGGGTTTAAACTCA |
| Insert R3 | AGAACCACCACCACCCGCATAATCCGGCACATCATACGGATAAGAACCACCACCACCAGA |
| Insert R4 | ACTAGCCTGCAGAAGCTTAAAAACATACTGTGTGTTTATGGGGCTGCCTTTGCTAGTCGTTGAGGG |
| Vector F | CATTTCTCCGTAAACAACATGTTTTCCCATGTAATATCCTTTTCTATTTTTCGTTCCG |
| Vector R | CTGCCTTTGCTAGTCGTTGAGGGGTGAGAACC |

**Protocol S1**. Conducting peptide library selection campaigns.

We have written this protocol to facilitate the use of the peptide library in selection campaigns, starting from the point of receiving the library from our commercial vendor (Kerafast, Cat# EF0023-FP) or via Material Transfer Agreement. We believe this protocol is accurate as of the publication date, but if any updates or revisions are necessary, the updated versions will be posted at https://github.com/kruselab/peptide-library.

### Protein reagent preparation

There are many reagent formats that are theoretically compatible with the use of this peptide library. We have found the most reliable success with biotinylated protein reagents and streptavidin-based secondary reagents, but other formats (e.g., directly labeling your protein with a fluorophore or using a labeled antibody directed against an epitope tag on your protein) may also work. We recommend preparing 20 nanomoles of your reagent and ensuring it is pure by SDS-PAGE and monodispersed by size exclusion chromatography before beginning a selection campaign.

### Thawing and expanding the peptide library

For most selections, starting from $10^{10}$ cells (which corresponds to sampling ~80% of total library diversity) will be sufficient to identify high-affinity binders and draw conclusions about motifs involved in binding interactions. In some situations, such as screening for short recognition motifs of 4–5 amino acids, substantially subsampling the library (e.g., using $10^7$ cells) may be adequate. In other situations, it may be desirable to thoroughly screen the library's diversity, which is possible but requires large culture volumes and multiple MACS columns. In this case, a number of cells 5–10 times the library diversity should be sampled (corresponding to 3–6 vials of $10^{10}$ cells). If this larger sampling rate is desired, everything in the following protocol should be scaled up by a factor of at least three (which ensures a >99% sampling of the library diversity).

1. Prepare 2 L of YGLC medium in a 3 L unbaffled Erlenmeyer flask.
2. Thaw the frozen vial containing $2 \times 10^{10}$ cells in a 30 °C water bath for 1–5 minutes.
3. With a sterile pipette, transfer the cells to the flask containing YGLC medium.
4. Grow the cells for 48 hours at 30 °C, shaking at ~200 rpm.
5. After 48 hours, remove 1 mL of culture and measure its $OD_{600}$. You may have to dilute the sample to remain in the linear range of your spectrometer. A fully saturated culture should have an $OD_{600}$ between 8–16. 1 AU corresponds to approximately $1.7 \times 10^7$ cells/mL.
6. Centrifuge the culture in sterile tubes (500 mL polypropylene tubes such as Corning Cat#431123 are convenient for this) at 3,500 *x g* for 5 minutes.
7. Discard the supernatants and resuspend the pellets in YGLC supplemented with 15% DMSO such that the final density is $5 \times 10^9$ cells/mL. The pellet will be large and its volume should be taken into account; thus, the final DMSO concentration should be close to 10%.
8. Prepare 2 mL aliquots in cryovials and place them in a cell freezing chamber. This protocol will yield about 18 vials, each containing $10^{10}$ cells.
9. Freeze at −80 °C for 48 hours.

### Performing selection campaigns

This protocol is intended to provide a starting point for selection campaigns using the peptide library, with the caveat that each target protein and campaign will have their own unique goals and requirements, and as such this protocol may have to be adjusted accordingly. The numbers provided here assume sampling $10^{10}$ cells, but if more thorough sampling of the library is desired, scale up the first MACS selection round by at least a factor of three (both for culture volumes and number of MACS columns used).

In general, a selection campaign will consist of multiple rounds of Magnetic-Activated Cell Sorting (MACS) and/or Fluorescence Activated Cell Sorting (FACS). MACS tends to be used in 1–2 early rounds to efficiently remove nonbinding clones, and FACS tends to be used in later rounds to more precisely control selection pressure. We recommend proceeding to FACS as early as possible; i.e., as soon as it becomes feasible to FACS sort ten times the number of cells eluted from the previous MACS round.

*MACS sorting*

Day 1: Start library

1. Thaw a vial of the library (or frozen previous selection round) in −trp +glu medium, at a density of ≤5 × $10^6$ cells/mL.
2. Grow the cells for 24–48 hours at 30 °C, shaking at ~200 rpm.

Day 2: Induce expression

3. Measure cell density as before, and passage a number of cells at least ten times the number of cells collected in the previous round (or $10^{10}$ cells for the first MACS round) into 2 L −trp +gal medium to induce peptide expression via the GAL promoter.
4. Grow the cells for 48 hours at 25 °C, shaking at ~200 rpm.

Day 4: MACS selection

*Negative selection (LD column)*

5. Measure density of –trp +gal culture as before.
6. Centrifuge [10 × previous library size] cells at 3,500 *x g* for 5 min at 4 °C. Discard supernatant.
7. Resuspend yeast pellet in 10 mL pre-chilled selection buffer.
8. Centrifuge at 3,500 *x g* for 5 min at 4 °C. Discard supernatant.
9. Resuspend pellet in 4.5 mL of selection buffer.
    - If using secondary reagents (e.g., a fluorophore-labeled antibody), add the secondary reagent at its working concentration, cover with foil, and incubate rotating for 20 min at 4 °C before proceeding.
10. Add 500 µL Miltenyi beads (often streptavidin or anti-biotin, although many different types are available for purchase) to yeast.
11. Incubate rotating for 40 min at 4 °C. After ~30 min, place LD column on magnetic rack with fins facing out. Equilibrate column with 5 mL selection buffer, keeping column covered to prevent contamination.
12. Centrifuge yeast at 3,500 *x g* for 5 min at 4 °C. Discard supernatant.
13. Resuspend in 5 mL selection buffer.
14. Add cells to column 1 mL at a time to prevent clumping, collecting flowthrough in a sterile 15 mL Falcon tube.
15. Discard LD column and plunger.

*Positive selection (LS column)*

16. Centrifuge LD column flowthrough at 3,500 *x g* for 5 min at 4 °C. Discard supernatant.
17. Resuspend pellet in 5 mL selection buffer containing antigen at its working concentration (often 1 µM).
    - If using secondary reagent, also add it at its working concentration now.
18. Incubate rotating for 1 hour at 4 °C.
19. Centrifuge at 3,500 *x g* for 5 min at 4 °C. Discard supernatant.
20. Resuspend in 4.5 mL selection buffer, then add 500 µL anti-fluorophore beads.
21. Cover with foil and incubate rotating for 20 min at 4 °C.
22. After ~15 min, place LS column on magnetic rack. Equilibrate column with 5 mL selection buffer.
23. Centrifuge at 3,500 *x g* for 5 min at 4 °C. Discard supernatant and resuspend in 5 mL of selection buffer.
24. Repeat wash and resuspend in 5 mL of selection buffer.
25. Remove 10 µL and keep on ice for later analysis. This will be the Load sample.
26. Flow yeast over column and collect flowthrough in a sterile 15 mL Falcon tube.
27. Wash column with 8 mL selection buffer. Record volume of flowthrough, then remove 10 µL and keep on ice. This will be the Flowthrough sample.
28. Remove column from magnet and hold over a sterile 15 mL Falcon tube.
29. Add 5 mL selection buffer to column and quickly use plunger to elute.

30. Remove 50 µL and keep on ice. This will be the Eluate sample.
31. Centrifuge eluate at 3,500 *x g* for 3 min at 4 °C. Discard supernatant. Pellet will be hard to see.
32. Resuspend in 3 mL –trp +glu medium and transfer to a 12 mL culture tube. Shake at 30 °C for 24 hours to recover.

*Measure recovery*

33. Measure yeast densities from Load, Flowthrough, and Eluate samples using hemocytometer or flow cytometer. The sum of cells in the Flowthrough and Eluate may not perfectly add up to the cells in the Load, but a successful selection Round 1 Eluate will typically contain between 1–50 × $10^6$ cells.

Day 5: Passage selected yeast into larger volume

34. Passage at least [10 × post-MACS library size] yeast into 20 mL –trp +glu.

Day 6: Passage into YGLC for storage

35. Passage [10 × post-MACS library size] into 20 mL YGLC media for freezing.

Day 8: Freeze aliquots

36. Spin YGLC culture (3,500 *x g*, 5 min) and discard supernatant.
37. Resuspend in YGLC + 10–15% DMSO (adjusting DMSO concentration such that final solution is 10%) at a density of ≤ 5 × $10^9$ cells/mL.
38. Aliquot at least [10 × post-MACS library size] cells into cryovials and freeze in cell freezing chamber.

If it is feasible to FACS sort [10 × post-MACS library size] cells, we recommend proceeding to FACS. If not, you can iteratively perform MACS, dropping the concentration of protein target each time to increase selection pressure.

*FACS sorting*

FACS sorting will depend much more than MACS on the specific goals of your campaign, but in general, you can follow the same Day 1 and Day 2 instructions under the MACS sorting section above to grow and induce the previous selection round. On Day 4, you can proceed to the following steps:

1. Measure cell density and pellet the required number of cells at 3,500 *x g* for 5 min. This is often one unstained sample ($10^7$ cells) and one sample stained with secondary reagents only ($10^7$ cells) that are used to set gates, and one sample with target protein and secondary reagents for sorting (ten times the previous round library size).
2. Resuspend in selection buffer at $10^7$ cells/mL. Meanwhile, prepare staining solutions and keep on ice.
3. Discard supernatants and resuspend pellets in staining solutions. Incubate rolling for 30 min at 4 °C.
4. Spin samples, discard supernatants, and resuspend in selection buffer.
5. Repeat wash and resuspend so final densities are $10^7$ cells/mL.
6. Sort sample into a tube containing 3 mL –trp +glu, using the unstained and secondaries-only samples to set appropriate gates (see Figure S1 for example gates). Transfer sorted sample into a 14 mL culture tube and grow shaking at 30 °C for 48 hours.

Following these steps, you can freeze this selection round as before starting from Day 5 under the MACS selection protocol above.

FACS can be repeated iteratively with different target protein concentrations, secondary reagents, or gating strategies until desired results are achieved.

**Buffer recipes**

**YGCL** (for amplifying libraries)
Add the following to a large beaker (dissolving requires vigorous stirring):

- 0.38% (3.8 g / L) -trp dropout media supplement
- 0.67% (6.7 g / L) yeast nitrogen base
- 2% (20 g /L) glucose
- 1.04% (10.4 g / L) sodium citrate
- 0.44% (4.4 g / L) citric acid monohydrate
- 1% (10 mL / L) pen/strep (10,000 units/mL stock)

Add DI water, adjust pH to 4.5, and sterile filter.

**-trp +glu/gal** (for selective growth (glucose) and induction (galactose)
Add the following to a large beaker (dissolving requires vigorous stirring):

- 0.38% (3.8 g / L) -trp dropout media supplement
- 0.67% (6.7 g / L) yeast nitrogen base
- 2% (20 g /L) glucose or galactose
- 1% (10 mL / L) pen/strep (10,000 units/mL stock)
- For plates, add 1.8% (18 g /L) bacto agar. Don't add glucose and pen/strep until after autoclaving.

Add DI water, adjust pH to 6.0, and sterile filter.

**Selection buffer***

- 20 mM HEPES pH 7.5
- 150 mM sodium chloride
- 0.1% (w/v) bovine serum albumin
- 5 mM maltose

Sterile filter before use.
*Many other buffers will work equally well—depending on your protein you may need to modify the composition.

**Catalog numbers**

LD columns (Miltenyi Biotec, Cat#130-042-901)
LS columns (Miltenyi Biotec, Cat#130-042-401)
Trp dropout (US Biological, Cat#D9531)
Yeast nitrogen base (Himedia, Cat#M878)
Glucose (Sigma, Cat#G8270)
Galactose (Difco, Cat#216310)
Pen/Strep (Gibco, Cat#15140-122)
Sodium citrate (Ward's Science, Cat#470302-530)
Citric acid (MP Biomedicals, Cat#150699)
